# Biosynthesis of Isonitrile Lipopeptides by Conserved Non-ribosomal Peptide Synthetase Gene Clusters in *Actinobacteria*

**DOI:** 10.1101/121228

**Authors:** Nicholas C. Harris, Michio Sato, Nicolaus A. Herman, Frederick Twigg, Wenlong Cai, Joyce Liu, Jordan Downey, Ryan Khalaf, Joelle Martin, Hiroyuki Koshino, Wenjun Zhang

## Abstract

A putative lipopeptide biosynthetic gene cluster is conserved in many species of *Actinobacteria*, including *Mycobacterium tuberculosis* and *M. marinum*, but the specific function of the encoding proteins has been elusive. Using both *in vivo* heterologous reconstitution and *in intro* biochemical analyses, we have revealed that the five encoding biosynthetic enzymes are capable of synthesizing a new family of isonitrile lipopeptides (INLPs) through a thio-template mechanism. The biosynthesis features the generation of isonitrile from a single precursor Gly promoted by a thioesterase and a non-heme iron(II)-dependent oxidase homologue, and the acylation of both amino groups of Lys by the same isonitrile acyl chain facilitated by a single condensation domain of a non-ribosomal peptide synthetase (NRPS). In addition, the deletion of INLP biosynthetic genes in *M. marinum* has decreased the intracellular metal concentration, suggesting the role of this biosynthetic gene cluster in metal transport.

**Significance Statement:** *Mycobacterium tuberculosis* is the leading causative agent of tuberculosis (TB), of which millions of deaths occur annually. A putative lipopeptide biosynthetic gene cluster has been shown to be essential for the survival of this pathogen in hosts, and homologous gene clusters have also been found in all pathogenic mycobacteria and other species of *Actinobacteria*. We have identified the function of these gene clusters in making a new family of isonitrile lipopeptides. The biosynthesis has several unique features, including an unprecedented mechanism for isonitrile synthesis. Our results have further suggested that these biosynthetic gene clusters play a role in metal transport, and thus have shed light on a new metal transport system that is crucial for virulence of pathogenic mycobacteria.

---

Small-molecule secondary metabolites are produced by microbes as chemical weapons to combat competing organisms, or as communication signals to control complex processes such as virulence, morphological differentiation, biofilm formation, and metal acquisition (1-3). One of the most important classes of secondary metabolites are non-ribosomal peptides, which are typically biosynthesized by modular non-ribosomal peptide synthetases (NRPSs) in an assembly-line manner (4). Two NRPS encoding gene clusters (*mbt* and *Rv0096-0101*) have been identified from the genome of *Mycobacterium tuberculosis*, the leading causative agent of tuberculosis (TB) that currently infects one-third of the world’s population. While the cluster of *mbt* has been characterized to biosynthesize mycobactin siderophores that form mycobactin-Fe(III) complexes for iron sequestration (5), the role of *Rv0096-0101* remains obscure despite various biological studies that have indicated the production of a virulence factor by this gene cluster (6-14). For example, using transposon site hybridization, *Rv0098* to *Rv0101* were predicted to be required for *M. tuberculosis* survival in a mouse model of infection (10). Consistently, a transposon insertion of *Rv0097* attenuated *M. tuberculosis* growth and survival in mice (7).

An *in silico* homology search has revealed that gene clusters homologous to *Rv0096-0101* are conserved in pathogenic mycobacteria, such as *M. bovis, M. leprae, M. marinum, M. ulcerans* and *M. abscessus* (**Fig. 1**), while absent in non-pathogenic mycobacteria such as *M. smegmatis*, providing further indication of the virulence-associated nature of the locus product in mycobacteria. Interestingly, in addition to the genus of *Mycobacterium*, related operons are found in the phylum of *Actinobacteria* across genera including *Streptomyces, Kutzneria, Nocardia*, and *Rhodococcus* (**Fig. 1**), suggesting a widespread presence of this cluster. Further bioinformatic analysis has shown that five genes (*Rv0097-0101*) are conserved across all identified gene clusters, and that these genes encode proteins homologous to an iron(II) and α-ketoglutarate (α-KG) dependent oxidase, a fatty acyl-CoA thioesterase, an acyl-acyl carrier protein ligase (AAL), an acyl carrier protein (ACP), and a single- or di-module NRPS, respectively (**Fig. 1 and S1**). Although all of these five proteins are typically involved in secondary metabolite biosynthesis, the identity of the corresponding metabolite and the specific function of these proteins have not yet been fully elucidated.

**Figure 1.**
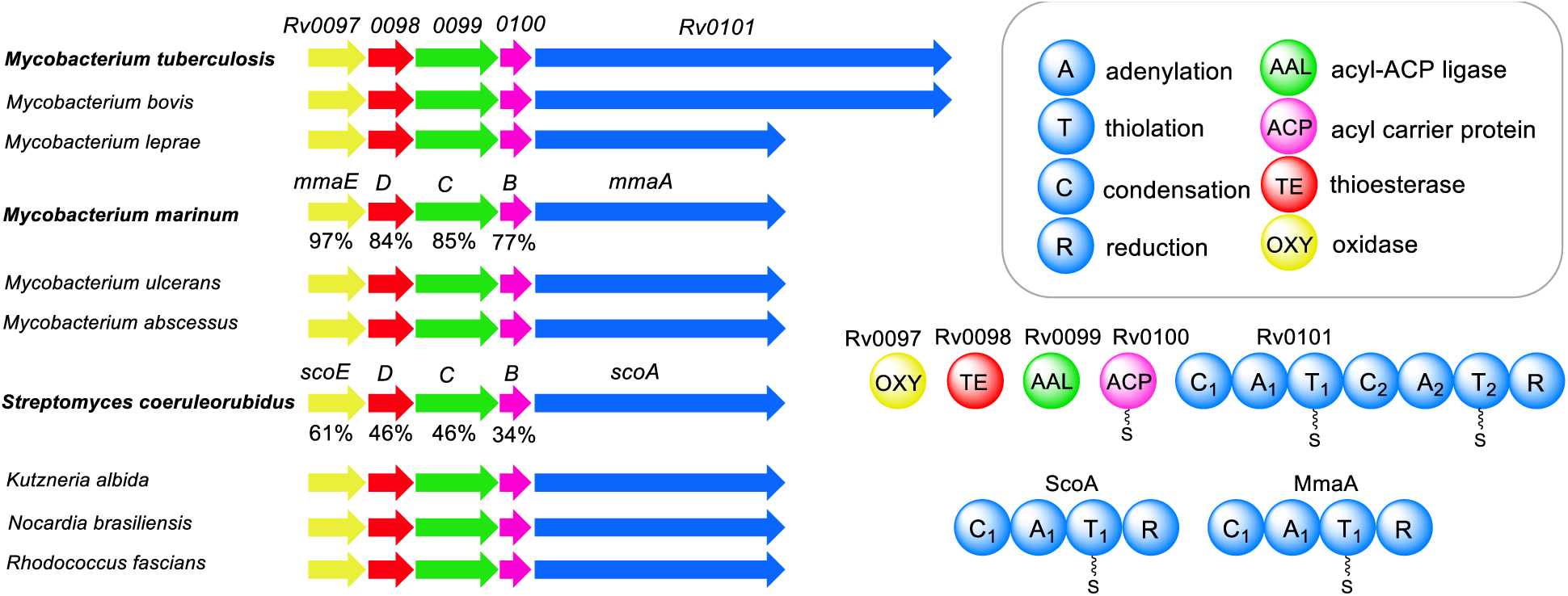
Schematic of the selected conserved biosynthetic gene clusters and their encoding protein products. Thousands of homologous gene clusters have been identified from published genomes. The similarity of protein homologues from *M. marinum* and *S. coeruleorubidus* to *M. tuberculosis* is shown below the gene clusters. A_1_ in Rv0101, MmaA, and ScoA was predicted to activate Lys based on the 10-residue specificity sequence (DIEDVGSVVK, DIEDVGSVVK, and DTEDVGTVVK, respectively). A_2_ in Rv0101 was predicted to activate Phe (DAWTVAAICK).

To better understand the role of this widespread gene cluster, we turned to reconstitution in *Escherichia coli* as a means to quickly and systematically assess the function of the five conserved enzymes through metabolomic exploration. The enzymes from *M. tuberculosis* H37Rv, *M. marinum* strain M (an opportunistic human pathogen), and *Streptomyces coeruleorubidus* NRRL18370 (15) (a known pacidamycin producer) were studied and compared to reveal similarities and variations in biosynthetic functions (**Fig. 1**). We discovered that these five conserved enzymes were necessary and sufficient to synthesize a unique group of isonitrile lipopeptides (INLPs). Based on both *in vivo* reconstitution and *in vitro* biochemical analysis, we scrutinized the timing and substrate specificity of these enzymes in INLP biosynthesis. Additionally, a mutagenesis study in *M. marinum* suggests that this biosynthetic gene cluster plays a role in metal acquisition.

## Results and Discussion

### Biosynthesis of INLPs by ScoA-E in *E. coli*

Previous biochemical studies of Rv0099-0101 suggested that a lipopeptide might be produced by these enzymes (16, 17). To unveil molecular features of the putative lipopeptide and gain preliminary knowledge of the functions of the five conserved biosynthetic enzymes, *scoA-E* from *S. coeruleorubidus* and *mmaA-E* from *M. marinum* (**Fig. 1**) were cloned for *E. coli* heterologous expression and functional reconstitution, respectively. A negative control strain transformed with empty vectors was also constructed. Protein expression analysis showed that all ten proteins were solubly expressed in *E. coli* (**Fig. S2**). The *E. coli* cultures were extracted with organic solvents, concentrated, and analyzed by Liquid Chromatography-High Resolution Mass Spectrometry (LC-HRMS) followed by untargeted metabolomics analysis using XCMS (18) for the determination of metabolic profile differences and the identification of new metabolites.

While no new metabolite was immediately identified in the culture of *E. coli-mmaA-E* compared to the negative control strain, upon co-expression of *scoA-E*, two new metabolites with molecular formulas C_18_H_28_N_4_O_4_ (**1**, calculated for C_18_H_29_N_4_O_4_^+^: 365.2183; found: 365.2185) and C_16_H_26_N_4_O_3_ (**2**, calculated for C_16_H_27_N_4_O_3_^+^: 323.2078; found: 323.2079) were found to be produced which were absent in the negative control (**Fig. 2** and **S3**). Both metabolites were UV-inactive above 210 nm and acid sensitive. The minor metabolite **2** was further predicted to be de-acetylated **1** based on HRMS/MS analysis (**Fig. S3**). To purify a sufficient quantity of **1** and **2** for structural elucidation, a total of ~40-L *E. coli* culture was prepared and extracted with chloroform, followed by purification via multiple rounds of HPLC using reverse-phase C18 columns. These purification steps yielded ~3 mg of pure compound **1** and ~0.5 mg of **2**. NMR spectra, including ^1^H, ^13^C, dqf-COSY, HSQC, HMBC, and ROESY spectra, were obtained for compound **1** and used to determine its molecular connectivity (**Fig. 2 and S4, Table S1**). The presence of an isonitrile moiety in **1** was further confirmed by comparison to the reported ^13^C-^14^N nuclear spin coupling constants and IR spectroscopy absorption (**Fig. 2**) (19). The absolute configuration of the C3’ and C3’’ (both R) was determined by acid hydrolysis to yield the monomer of 3-amino butyric acid which was then reacted with Marfey’s reagent and compared to the standards (**Fig. 2**) (20). The NMR spectra of **2** confirmed that it is a C1 de-acetylated analogue of **1 (Fig. 2 and S5, Table S2**). The molecular structures of **1** and **2** were revealed to be similar to two known isonitrile antibiotics (SF2768 and SF2369) that were originally isolated from the culture filtrates of different *Actinomycetes* species (21-23) (**Fig. 2**), strongly indicating that *E. coli* was a suitable heterologous host for the reconstitution of ScoA-E activities to produce relevant metabolites. Systematic removal of each of the five genes from *E. coli-scoA-E* abolished the production of **1** and **2**, demonstrating the necessity of all five enzymes in synthesizing these two INLPs (**Fig. 2**).

**Figure 2.**
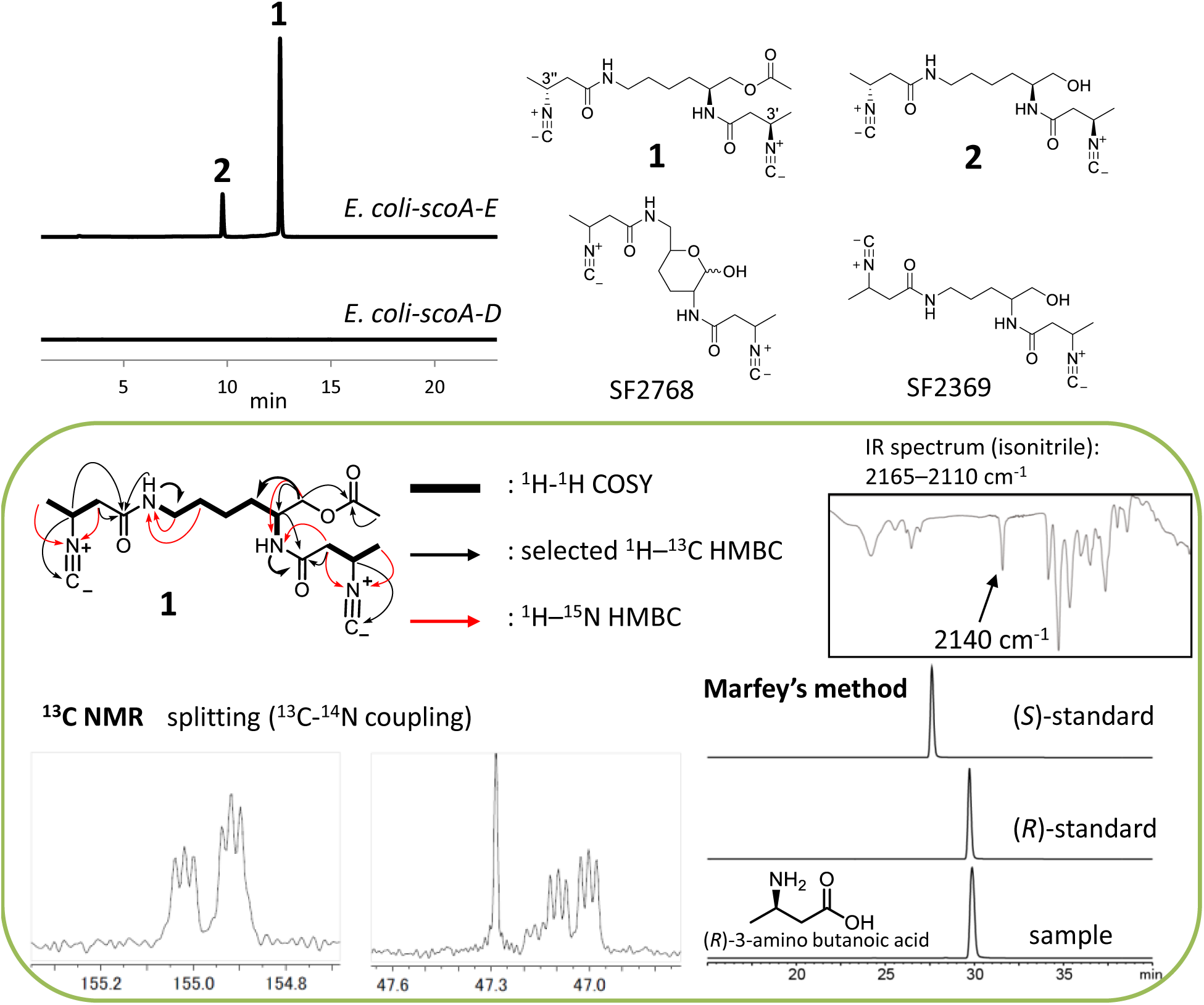
Biosynthesis of INLPs (1 and 2) by ScoA-E in *E. coli*. Extracted ion chromatograms show production of **1** and **2** by *E. coli-scoA-E*. The calculated masses for **1** and **2** with 10 ppm mass error tolerance were used for each trace. The negative control strain with empty vectors and strains containing any of the four gene combination did not produce **1** and **2**, and only one representative trace is shown here for simplicity. The structural determination of **1** is boxed. Structures of two known *Actinomycetes* metabolites are also shown here.

### Proposed Biosynthetic Pathways for INLPs by the Five Conserved Biosynthetic Enzymes

The heterologous production of **1** and **2** in *E. coli* enabled the assignment of function to the five conserved biosynthetic enzymes, especially the role of the two modification enzymes (ScoD and ScoE), in a unique isonitrile moiety synthesis. We have thus proposed putative enzymatic pathways for INLP biosynthesis (**Fig. 3**). **2** is presumably generated through a hybrid AAL-NRPS assembly line-based mechanism. The assembly line starts with the activation of crotonic acid by ScoC through adenylation, and the adenylated acid is loaded onto an ACP (ScoB) for further processing. The α,β-unsaturated fatty acyl-ACP is then modified by a thioesterase homologue (ScoD) and a non-heme iron(II)-dependent oxidase (ScoE) to generate a β-isonitrile fatty acyl-ACP intermediate. Instead of being a thioesterase as suggested by the previous biochemical and structural analyses of Rv0098 (24), we propose ScoD to be a reverse lyase-like enzyme that catalyzes a Michael addition of Gly to the β-position of an α,β-unsaturated fatty acyl-ACP to yield an *N*-carboxymethyl-3-aminoacyl-ACP. ScoE presumably catalyzes the subsequent oxidation and decarboxylation to yield a β-isonitrile moiety. The isonitrile-modified fatty acyl chain is then condensed to both amino groups of Lys promoted by the NRPS (ScoA) and reductively released by the reduction (R) domain of the NRPS to form a terminal alcohol product **2**. This reductive release mechanism is consistent with the identified activity of the R domain in Rv0101, which contains a conserved Ser/Tyr/Lys catalytic triad and was demonstrated to catalyze a four-electron thioester reduction using the purified truncated R domain and a synthetic substrate (17). We propose that the extra acetyl moiety found in **1** is due to the activity of a promiscuous acetyltransferase endogenous to *E. coli* for possible detoxification, and similar acetylation events have been observed in other shunt product biosynthesis (25). An analogous biosynthetic pathway involving MmaA-E has also been proposed, differing in the chain length of the fatty acyl moiety (**Fig. 3**). Although ScoC seems to prefer a short-chain fatty acid substrate (such as crotonic acid as shown in **1** and **2**), MmaC likely favors a medium-chain fatty acid substrate (such as 2-decenoic acid) since previous biochemical and structural analyses of Rv0099, a homologue of MmaC from the same genus of *Mycobacterium*, showed that this AAL activated fatty acids of medium-chain length (16, 17).

**Figure 3.**
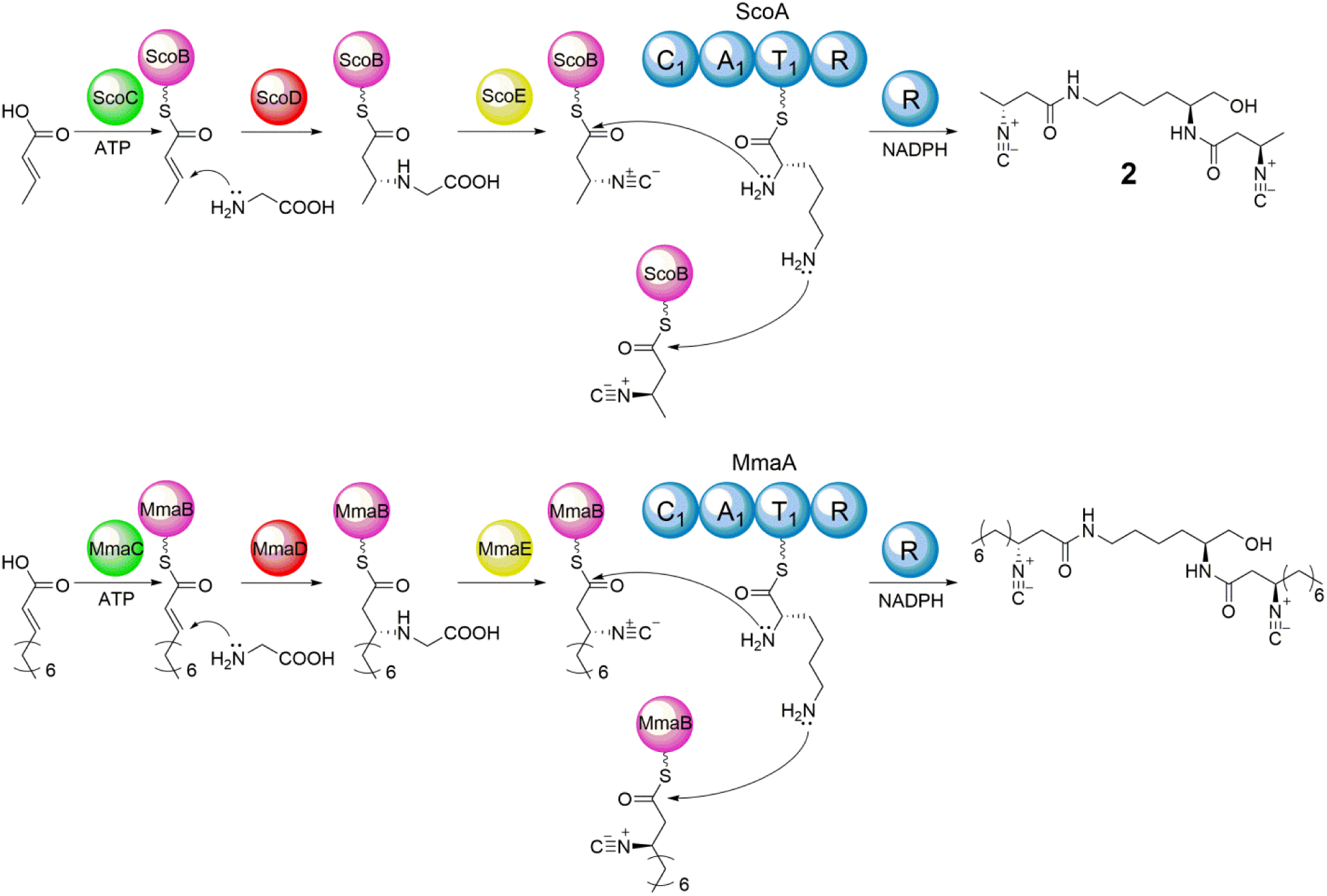
Proposed function of ScoA-E and MmaA-E in INLP biosynthesis.

### AAL, ACP, and NRPS Promote Di-acylated Lipopeptide Biosynthesis

We next performed additional *in vivo* metabolic analyses using various combinations of biosynthetic genes and *in vitro* biochemical analyses using purified enzymes to dissect the proposed biosynthetic pathways for INLPs (**Fig. 3**). The substrate specificity of NRPSs, including ScoA and MmaA, was tested using the classical ATP-[^32^P]PPi exchange assay. As expected, both enzymes demonstrated a strong preference for the activation of L-Lys (**Fig. 4a**), which is consistent with the molecular structures of **1** and **2** and supports the assignment of the absolute configuration of C2 in **1** to be *2S* (**Fig. 2**). Although L-Ornithine was not activated by ScoA/MmaA, it could be a preferred substrate for other conserved NRPSs based on the structure of SF2369 (**Fig. 2**). We next probed the fatty acid substrate specificity of AALs encoded by *scoC* and *mmaC*, respectively. The ability of ScoC to reversibly adenylate various acids was tested using the ATP-[^32^P]PPi exchange assay. ScoC exhibited a strong preference for the activation of fatty acids with a short-chain length (C4-C8), and as expected, α,β-unsaturated fatty acids were well recognized (**Fig. 4a**). The subsequent loading of selected fatty acyl moieties onto ScoB was further confirmed by HRMS analysis (**Fig. 4b**). MmaC demonstrated an intrinsic ATPase activity in the ATP-PPi exchange assay which prohibited the determination of substrate specificity using this method. Nonetheless, the direct substrate activation and loading assays confirmed that similar to Rv0099, MmaC preferentially activates fatty acids of medium-chain length with tolerance toward α,β-unsaturation (**Fig. 4b**).

**Figure 4.**
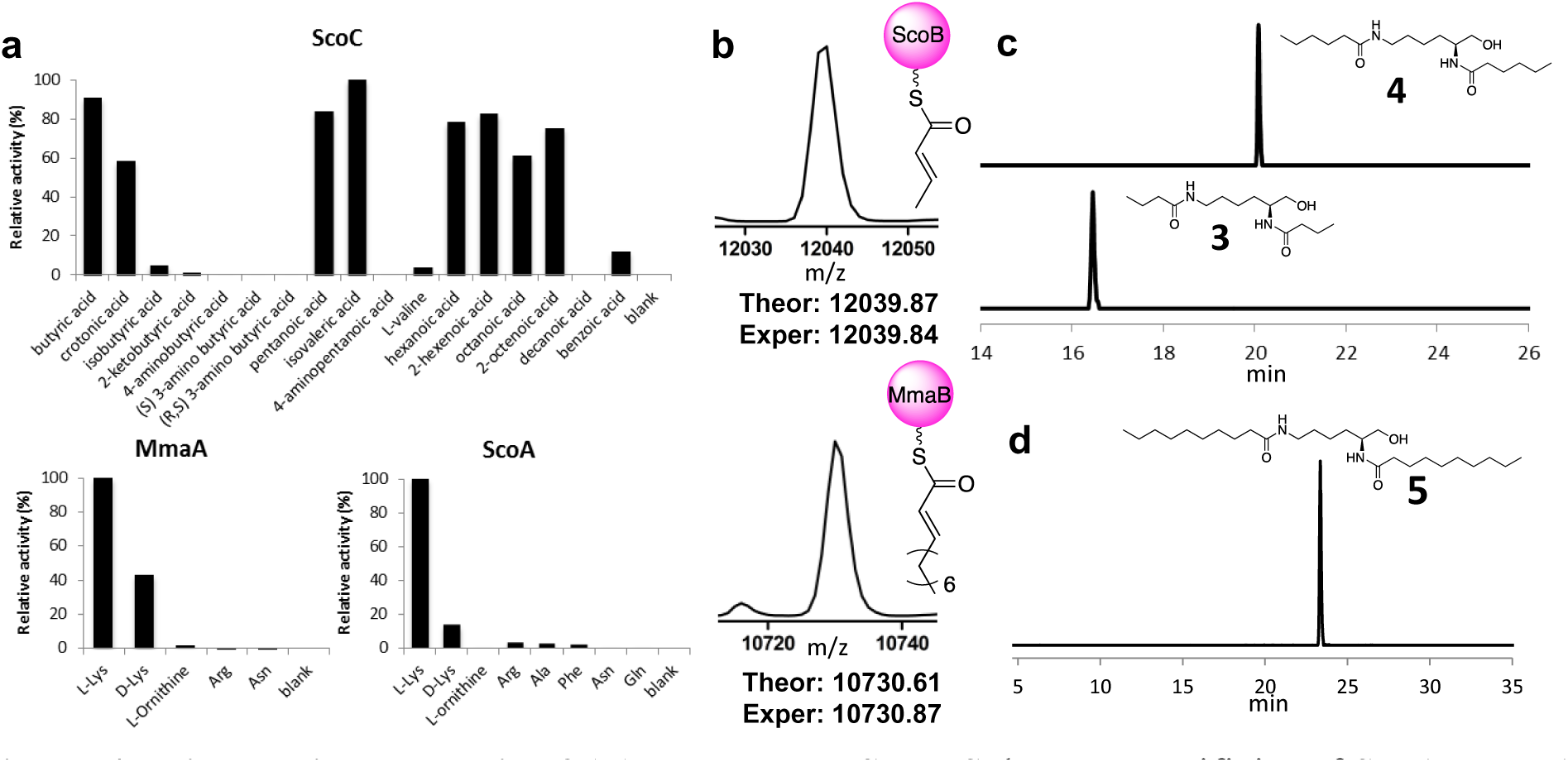
Biochemical analysis of AALs and NRPSs. **a**, Substrate specificity of ScoA, MmaA, and ScoC determined by ATP-[^32^P]PPi exchange assays. **b**, Detection of crotonyl-S-ScoB and decenoyl-S-MmaB by MS with maximum entropy deconvolution. **c**, Extracted ion chromatograms showing the production of **3** and **4** in ScoA-C assays using butyric acid and hexanoic acid as a substrate, respectively. **d**, Extracted ion chromatogram showing the production of **5** in the MmaA-C assay using decanoic acid as a substrate. The calculated masses for **3-5** with 10 ppm mass error tolerance were used for each trace.

One of the unusual events in the proposed biosynthetic pathway is the condensation of the same fatty acyl moiety to both amino groups of Lys, presumably promoted by a single C domain of the NRPS (**Fig. 3**). We hypothesized that this C domain has relaxed substrate specificity, and the product formation assays using ScoA-C or MmaA-C successfully yielded lipopeptides (**3-5**) with the amide bond formation at both amino positions (**Fig. 4c, 4d, and S6**). Negative controls missing any of the enzyme or substrate (fatty acid, Lys, ATP, and NADPH) abolished the production of **3-5**. Products with a terminal alcohol were generated in these assays, confirming the four-electron reduction activity of the conserved R domain in ScoA and MmaA.

### Homologues of Rv0097 and Rv0098 Promote Isonitrile Biosynthesis through an Unprecedented Mechanism

Although the isonitrile functionality has been found in quite a few natural products, only one biosynthetic pathway has been identified in which one carbon is transferred to an amino group catalyzed by an isonitrile synthase such as IsnA (26-29). Our proposed pathway for isonitrile synthesis is mechanistically distinct from the known pathway by using a different set of enzymes. Rv0098 has previously been shown to be a long-chain fatty acyl-CoA thioesterase, although structurally, it lacks a general base or a nucleophile that is conserved in the thioesterase catalytic site. In addition, very low hydrolysis activities were obtained in the biochemical analysis of Rv0098, which solicited further detailed characterization of this hypothetical protein (24). The recent biochemical characterization of CmiS1, a homologue of ScoD (identity/similarity = 47%/56%), showed that CmiS1 catalyzed the Michael addition of Gly to the β-position of a non-2-enoic acid thioester in the biosynthesis of the macrolactam antibiotic cremimycin (30). We thus have proposed that a similar reaction of Michael addition of Gly to the β-position of an α,β-unsaturated fatty acyl-ACP could be promoted by ScoD/MmaD/Rv0098 to yield an *N*-carboxymethyl-3-aminoacyl-ACP (**Fig. 3**). This is consistent with the failed activation of 3-amino butyric acid by ScoC **(Fig. 4a**), which argued against the known pathway using an isonitrile synthase. To confirm the proposed function of MmaD, a biochemical reaction using MmaB-D, ATP, 2-decenoic acid, and Gly was performed, and the product was analyzed by LC-HRMS after release from the protein by base hydrolysis. This yielded the expected Gly adduct (calculated for C_12_H_24_NO_4_^+^: 246.1700; found: 246.1701), and its mass was shifted by +1 using [2-^13^C]Gly, or by +2 using [2-^13^C, ^15^N]Gly as an alternative substrate (**Fig. 5a and S7**). Negative controls omitting any of the three proteins or substrate abolished the production of the Gly adduct, indicating that the formation of the Gly adduct requires the activation of the fatty acid substrate and occurs on a thio-templated assembly line (**Fig. 5a**). Similar reactions were also performed using ScoB-D and the activity of ScoD in forming a Gly adduct was also confirmed **(Fig. S8**). We further observed that the Gly adduct on ACP (either MmaB or ScoB) was readily released by hydrolysis *in vitro*, likely promoted by MmaD/ScoD, consistent with the observed thioesterase activity of Rv0098 and CmiS1 (5, 18). We thus could not reconstitute the activity of MmaE/ScoE, the iron(II) and α-KG dependent oxidase homologue, in promoting the subsequent oxidation and decarboxylation *in vitro*, most likely due to substrate limitation.

**Figure 5.**
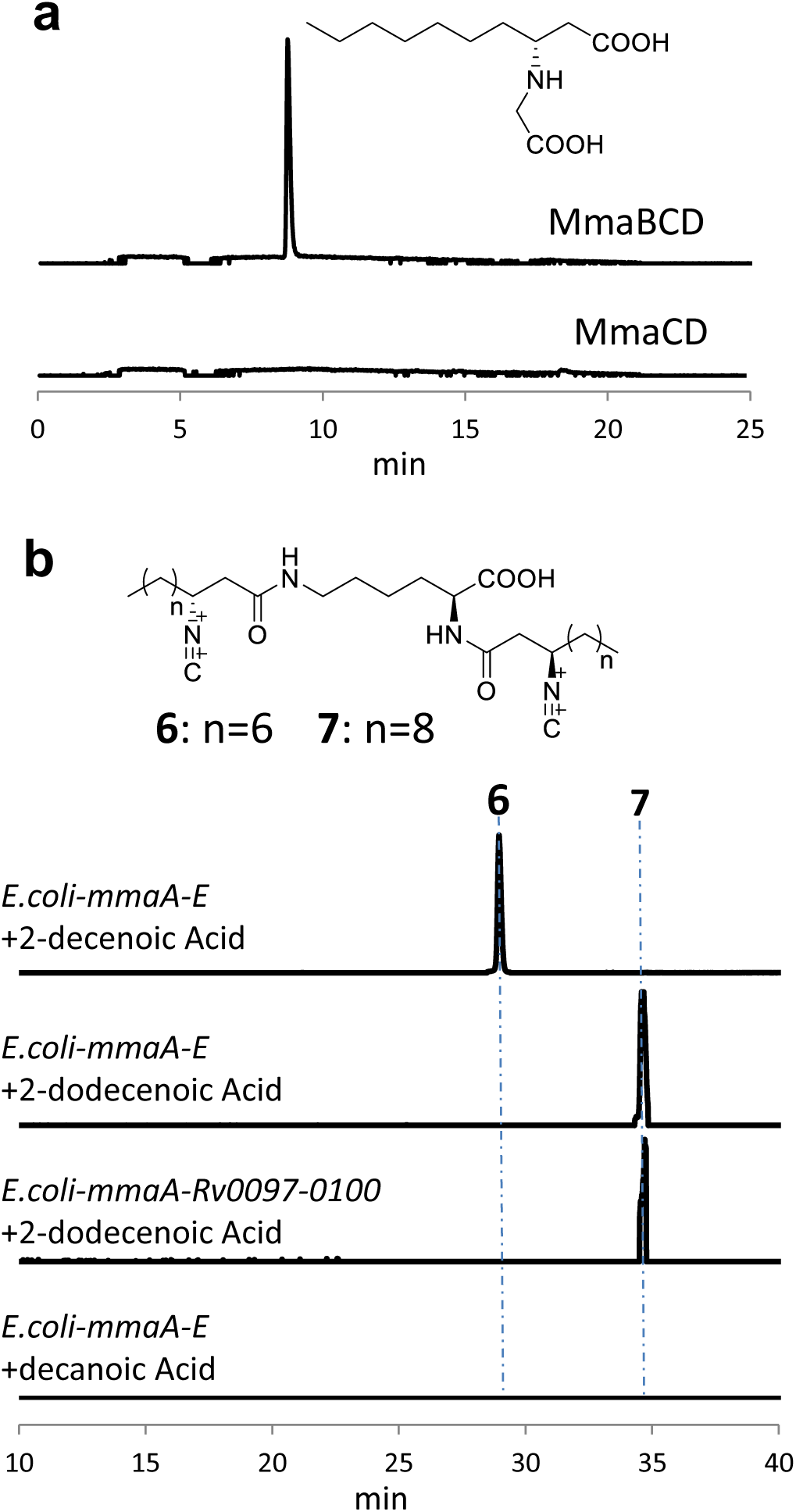
*In vitro* and *in vivo* analysis of isonitrile formation. **a**, Extracted ion chromatogram showing the production of Gly adduct in the MmaB-D assay. The calculated mass with 10 ppm mass error tolerance was used for each trace. Control assays missing any of the protein or substrate (ATP, 2-decenoic acid, and Gly) abolished the production of Gly adduct, and only one representative trace with no MmaB is shown for simplicity. **b**, Extracted ion chromatograms showing the *E*. coli-based production of **6** and **7** after feeding of 2-decenoic acid and 2-dodecenoic acid, respectively. The calculated masses for **6** and **7** with 10 ppm mass error tolerance were used for each trace. Strains containing any of the four gene combination, or feeding of decanoic acid and dodecanoic acid, did not produce **6** and **7**, and only one representative trace with the feeding of decanoic acid to *E. coli-mmaA-E* is shown here for simplicity.

Since reconstituting the activity of MmaE/ScoE *in vitro* has not yet been successful, we turned to *in vivo* reconstitution coupled with feeding of isotope labeled Gly to further confirm the proposed activity of enzymes in isonitrile biosynthesis. While co-expression of *scoA-C* in *E. coli* yielded expected di-acylated lipopeptides such as **3** and acetylated **3**, the addition of *scoD* or *scoE* to *E. coli-scoA-C* did not change the metabolic profile with no new metabolite identified through comparative metabolomics, indicating that the modification of the fatty acyl chain occurs before the biosynthetic intermediate release by the NRPS, and the C or R domain of ScoA is intolerant of a bulky side group on the fatty acyl chain (Gly adduct). Only upon the addition of both *scoD* and *scoE* to *E. coli-scoA-C*, new major products of **1** and **2** were identified (Fig. 2a). We thus reasoned that ScoD and ScoE function on the assembly line, most likely on the free standing ACP, ScoB, for isonitrile synthesis. Additionally, feeding of 10 mM of Gly to the culture of *E. coli-scoA-E* boosted the titer of **1** by over 20-fold, and feeding of [2-^13^C, ^15^N]Gly showed that C(2)-N of Gly was incorporated into the isonitrile group of **1** and **2**, supporting the proposed role of ScoD and ScoE in isonitrile biosynthesis **(Fig. S3**).

To further confirm the role of MmaA-E in INLP biosynthesis, we reattempted the *in vivo* reconstitution of activities of MmaA-E in *E. coli*. Based on the substrate specificity of MmaC, we reasoned that the *in vivo* substrate limitation could be the reason for the failed INLP production by *E. coli-mmaA-E*. 2-decenoic acid or 2-dodecenoic acid was then fed to the culture of *E. coli-mmaA-E* and untargeted metabolomics analysis was performed to search for new metabolites. This led to the identification of a trace amount of two new metabolites with molecular formulas C_28_H_48_N_4_O_4_ (**6**, calculated for C_28_H_49_N_4_O_4_^+^: 505.3748; found: 505.3748) and C_32_H_56_N_4_O_4_ (**7**, calculated for C_32_H_57_N_4_O_4_^+^: 561.4374; found: 561.4376), respectively (**Fig. 5b and S9**). HRMS analysis suggested that **6** and **7** are INLPs similar to **1** and **2**, but contain two longer fatty acyl chains and a C1 acid moiety **(Fig. S9**). The presence of the isonitrile was further confirmed by IR spectroscopy to show signature absorption of 2132 cm^−1^. As expected, the production of **6** and **7** was dependent on the co-expression of all five genes of *mmaA-E*, and feeding of [2-^13^C, ^15^N]Gly to the culture of *E. coli-mmaA-E* demonstrated that C(2)-N of Gly was incorporated into **6 (Fig. 5b and S9**). It is notable that feeding of decanoic acid or dodecanoic acid did not lead to the production of **6** and **7**, supporting our hypothesis that the biosynthesis of INLPs requires the activation of an α,β-unsaturated fatty acid (**Fig. 3**). In addition, replacement of *mmaB-E* by *Rv0097-0100* in *E. coli* produced **7** upon the feeding of 2-dodecenoic acid, demonstrating the interchangeable property of the encoding four enzymes (**Fig. 5b**). The INLP alcohols could not be detected, suggesting that 2-decenoic acid and 2-dodecenoic acid may not be the native substrate for the *M. marinum* proteins, which led to the impaired activity of AAL-NRPS assembly line and low titers of the corresponding INLPs in *E. coli*. Indeed bioinformatic analysis showed that putative fatty acid modification enzymes are encoded in close proximity to *mmaA-E* in the *M. marinum* genome **(Fig. S1**).

### Implications for the Role of the Conserved Gene Clusters in Metal Transport

The isonitrile functionality is known to behave as an electron-rich analogue of carbon monoxide and form coordination complexes with most transition metals (31, 32). Metals are required in many life processes, and pathogenic bacteria have addressed the challenge of deficiencies in essential metals or high concentrations of toxic metal cations in the host by evolving metal homeostasis mechanisms that are frequently associated with virulence (33-40). The operon of *Rv0096-0101* has been known to be critical for the virulence of *M. tuberculosis* and essential for the survival of this pathogen in macrophages and mice (7, 9, 10, 26, 31, 32). Based on the metal binding ability of isonitrile, we postulated that this conserved biosynthetic gene cluster plays a role in metal transport and homeostasis. *In silico* analysis demonstrated that putative heavy metal translocating P-type ATPases such as CtpA and CtpB are encoded in close proximity to the INLP biosynthetic genes in the *M. tuberculosis and M. marinum* genomes **(Fig. S1**), and the expression of CtpB was shown to be co-regulated with *Rv0096-0101* in *M. tuberculosis* by transcriptomic analysis (41). In addition, it has been reported that the expression of the *Rv0096-0101* gene cluster was normally repressed in many synthetic media including 7H9 where metal concentrations are high, but was constantly induced in Sauton’s medium where metal concentrations are relatively low (6), suggesting the metal uptake function of the cluster. We next constructed a mutant strain of *M. marinum* devoid of *mmaA-E* through allelic exchange according to the reported method (42), and the intracellular metal content of the *M. marinum* wild-type and Δ*mmaA-E* was determined to probe the function of these genes. Deletion of the gene cluster caused a significant decrease in the intracellular accumulation of zinc in Sauton’s medium as shown by Inductively Coupled Plasma-Mass Spectrometry (ICP-MS) analysis (**Fig. 6**). This is consistent with the finding from microarray analysis that the transcription of *Rv0097* was downregulated in *M. tuberculosis* exposed to excess zinc (37). These results strongly indicate that the conserved gene clusters synthesize a unique family of INLPs that promote metal transport in pathogenic mycobacteria.

**Figure 6.**
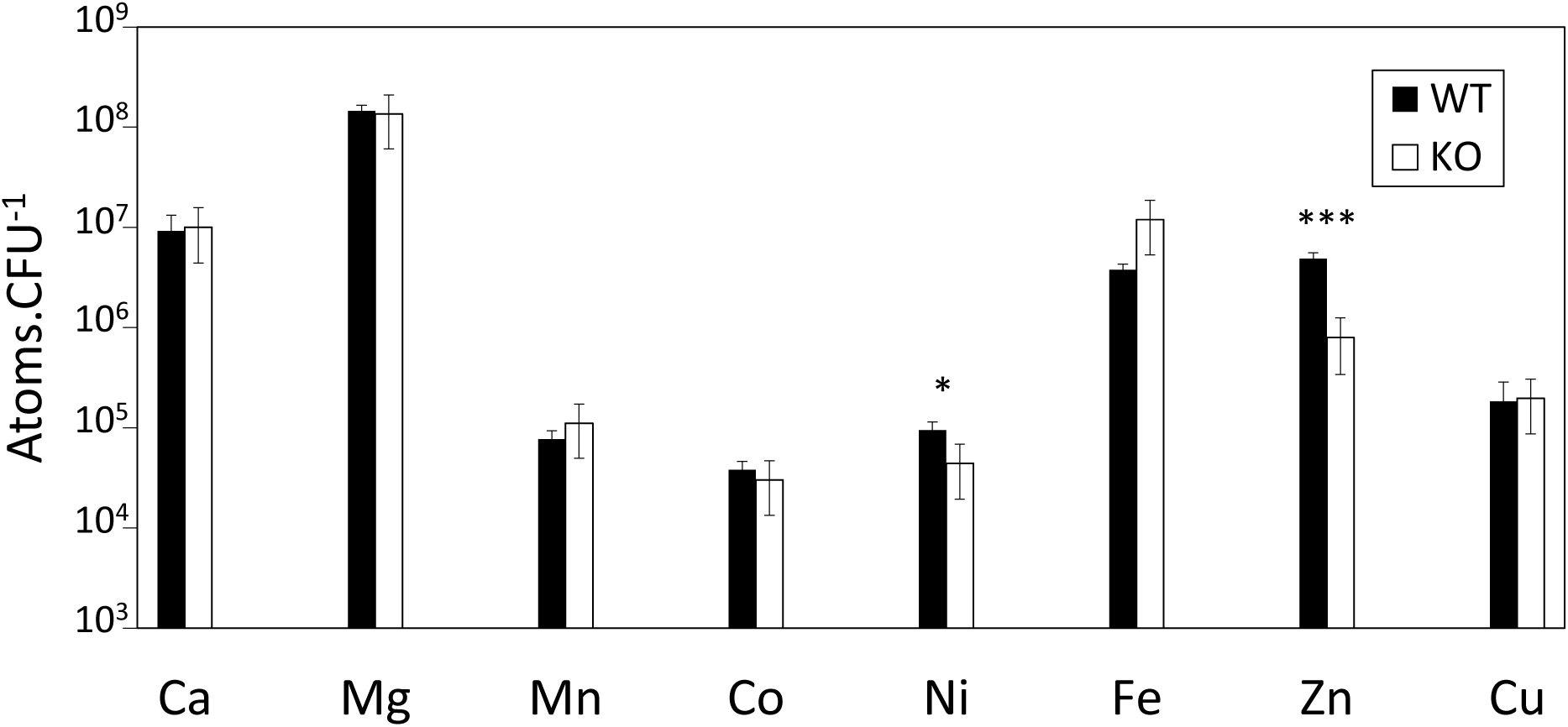
Effect of *mmaA-E* mutation in metal content. Intracellular metal content of the *M. marinum* wild-type (left) and Δ*mmaA-E* (right) strains grown in Sauton’s medium was determined by ICP-MS. Error bars, mean ±SD *P ≤ 0.05 and *** *P* ≤ 0.001.

## Conclusions

We have revealed the function of a biosynthetic gene cluster that is widespread in *Actinobacteria* including pathogenic mycobacteria such as *M. tuberculosis* and *M. marinum*. Using both *in vivo* and *in vitro* analyses, we discovered that the conserved five biosynthetic enzymes were capable of synthesizing a new family of INLPs by using a thio-template mechanism. This unusual biosynthetic pathway starts with the activation and loading of an α,β-unsaturated fatty acid onto an ACP by a promiscuous AAL. A Michael addition of Gly to the β-position of the α,β-unsaturated fatty acyl-ACP is then promoted by a thioesterase homologue, followed by oxidation and decarboxylation, presumably catalyzed by a non-heme iron(II)-dependent oxidase, to generate a β-isonitrile fatty acyl-ACP intermediate. This isonitrile intermediate is then condensed to both amino groups of Lys promoted by an NRPS and reductively released to form a terminal alcohol product. The identified isonitrile biosynthetic pathway is distinct from the canonical mechanism in which one carbon is transferred to an amino group to form isonitrile catalyzed by an isonitrile synthase (26-29). In addition, the condensation of an acyl moiety to both amino groups of Lys, presumably promoted by a single C domain of the NRPS, is rare in non-ribosomal peptide biosynthesis. Further mutagenesis study in *M. marinum* suggests that this biosynthetic gene cluster plays a role in metal transport. This study thus sheds light on a new metal transport system that is critical for the virulence of pathogenic mycobacteria. Future study of native INLP metallophores in mycobacteria and their mode of action during infection may inspire new therapies to combat pathogenic mycobacteria, in particular *M. tuberculosis*, from which millions of people die every year.

## Materials and Methods

Full experimental details are available in SI Appendix, Materials and Methods.

## Acknowledgement

We thank M. Hadjithomas (JGI) for assistance with bioinformatics search of homologous gene clusters, J. Pelton (UC Berkeley) for assistance with NMR spectroscopic analysis, J. Cox (UC Berkeley) for providing plasmids for *M. marinum* genetic manipulation, M. Welch (UC Berkeley) for providing *M. marinum* strain M, W. Yang (UC Berkeley) for assisting with ICP-MS analysis, S. Stanley and K. Sogi (UC Berkeley) for providing assistance with *M. marinum* genetic manipulation, and S. Bauer (UC Berkeley) for assisting with LCMS analysis. N. Harris is funded by the National Science Foundation Graduate Research Fellowship Program. This research was financially supported by the Pew Scholars Program and the National Institutes of Health (DP2AT009148).

## Author Contributions

N. C. H. and W. Z. designed the experiments, analyzed the data and wrote the manuscript. N. C. H, W. Z., J. L., N. A. H., and J. D. constructed plasmids. N. A. H. and J. L. performed comparative metabolomics experiments. M. S. performed purification of isonitrile lipopeptides. M. S., H. K., and W. C. performed NMR experiments and structure characterization. N. C. H., M. S., J. L., J. D., R. K., and J. M., purified proteins and performed biochemical assays. N. C. H. and W. C. performed MS/MS experiments. N. C. H. and F. T. performed mutagenesis and ICP-MS experiment. All authors discussed the results and commented on the manuscript. This project is supervised by W. Z.

